# Ubiquitylated H2A.Z is associated with diverse types of silenced chromatin including methylated CpG islands and homopurine/homopyrimidine sequences

**DOI:** 10.1101/759852

**Authors:** Bakhtiyar Taghizada, Marlee K. Ng, Ulrich Braunschweig, Benjamin J. Blencowe, Peter Cheung

## Abstract

H2A.Z monoubiquitylation has been linked to transcriptional repression, but the molecular details involved remain unclear. To address this, we developed a unique ubiquitylation-dependent auto-biotinylation approach, named H2A.Z-UAB, to purify ubiquitylated H2A.Z (H2A.Zub) mononucleosomes for biochemical and genome-wide analyses. We found that H2A.Zub nucleosomes are enriched for the repressive histone post-translational modification H3K27me3 but depleted of H3K4 methylation and other active transcription-associated modifications. ChIP-seq analyses revealed that H2A.Zub-nucleosomes associate with repressive chromatin states and are mostly enriched at genes with low or no expression. Consistent with the biochemical analyses, H2A.Zub ChIP-seq peaks align with regions depleted of H3K4 methylation, H3K27 acetylation, but enriched for H3K27me3. We further observed a strong correlation between high levels of H2A.Zub and DNA methylation, particularly at methylated CpG-rich islands. Finally, we found that a significant proportion of H2A.Zub peaks correspond to homopurine/homopyrimidine sequences that are generally linked to silenced genes and repressive chromatin. Collectively, these findings suggest that H2A.Z ubiquitylation plays an overarching role in global transcriptional silencing through its association with multiple types of repressive mechanisms.

**GRAPHICAL ABSTRACT:** 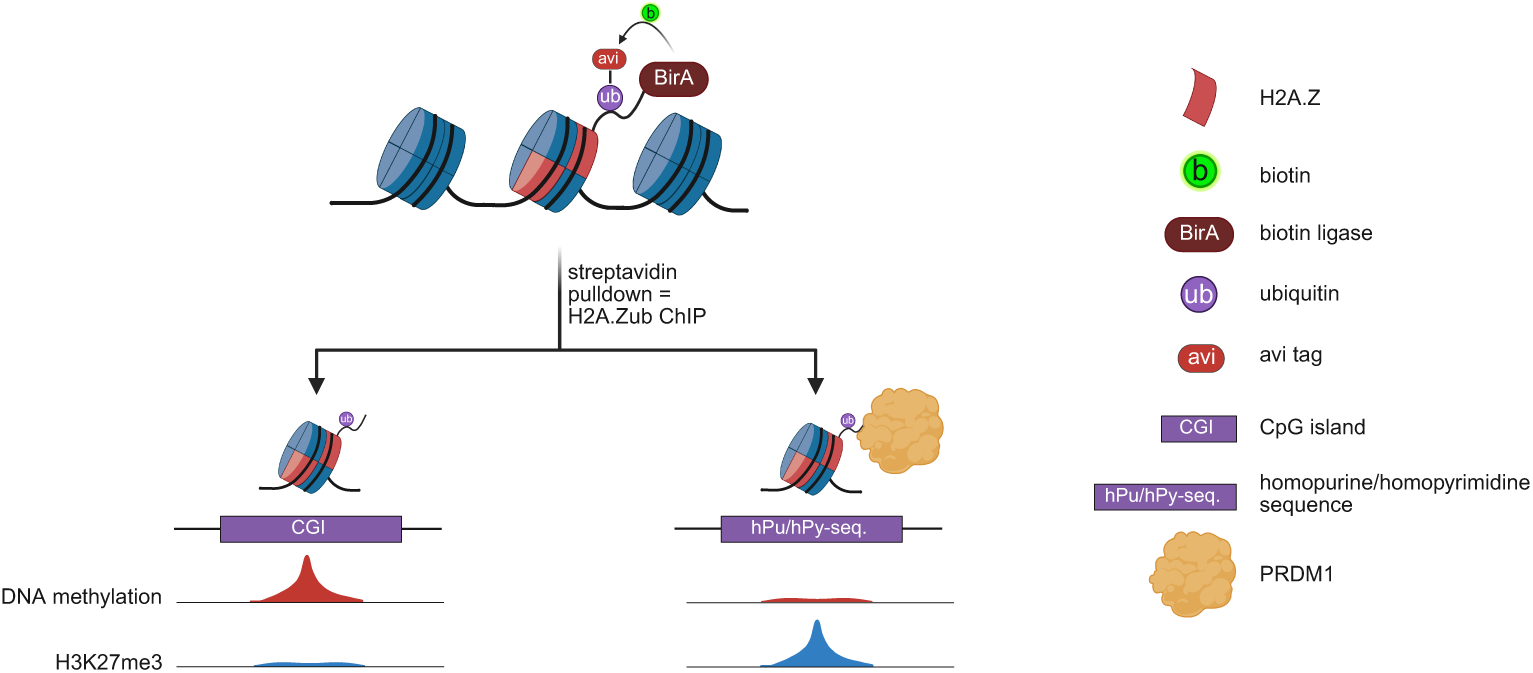

## INTRODUCTION

Fine-tuning of chromatin organization is paramount for regulating genome accessibility and orchestrating a myriad of DNA-templated processes such as transcription and DNA repair. The basic unit of chromatin is the nucleosome, a nucleoprotein complex made up of ∼ 147 bp of DNA wrapped around a histone octamer comprising 2 copies each of the core histones H2A, H2B, H3 and H4 (1). The addition of various histone post-translational modifications (PTMs) and the substitution of core histones with their non-canonical isoforms (i.e., histone variants) further confer distinct properties and functions to individual nucleosomes, contributing to dynamic organization and regulation of chromatin states and of the entire genome (2).

H2A.Z is a variant within the H2A family that replaces the canonical histone at strategic genomic regions (3–6). In vertebrates, this variant has two main isoforms (H2A.Z.1 and H2A.Z.2), encoded by separate genes, along with a splice variant of H2A.Z.2 known as H2A.Z.2.2 (7–9). In contrast, other organisms typically have a single version of H2A.Z (10). Knockout of H2A.Z.1 in mice or H2A.Z in other organisms is generally lethal, though exceptions exist (e.g., in *S. cerevisiae*) (11–13). To date, the functions of H2A.Z that are essential for viability are incompletely understood (14,15). H2A.Z is enriched at nucleosomes flanking transcription start sites (TSSs) of numerous genes, as well as at regulatory regions such as enhancers and promoters (16–19). This distribution aligns with a wealth of evidence linking H2A.Z to transcriptional regulation (15). Knock-down and functional studies have shown that H2A.Z is linked to both transcriptional activation and repression, with these contrasting activities likely mediated through its differential PTMs (14,15,20). Acetylation of H2A.Z directly correlates with transcriptional activation, whereas monoubiquitylation at its C-terminus is associated with transcriptional repression (3,6,21–24).

C-terminal monoubiquitylation of H2A.Z (H2A.Zub), was originally found to be mediated by the RING1B E3 ligase of Polycomb Repressor Complex 1 (PRC1), and enriched on the inactive X chromosome in human female cells (23). Later studies identified this modified H2A.Z at the promoters of androgen receptor-regulated genes in prostate cancer cells and at bivalent promoters of murine embryonic stem cells (mESCs), where it imparted transcriptionally repressive functions (5,22,24). At bivalent promoters of mESCs, H2A.Zub promotes transcriptional silencing in part by antagonizing recruitment of the transcriptional activator Brd2 (24). However, thorough analysis of H2A.Zub in the genome-wide context, and how it associates with other epigenetic features or mechanisms remain insufficiently elucidated.

Comprehensive analysis of the functional role of H2A.Z ubiquitylation has been hindered by the lack of specific reagents or high-quality antibodies against this form of H2A.Z. An earlier study reported the inability to obtain high-quality ChIP-seq data using an H2A.Zub antibody, while another study relied on the cross-reactivity of an H2AK119ub antibody towards H2A.Zub (5,24). To address this limitation and to facilitate mechanistic studies of H2A.Zub, we have developed a novel non-antibody-dependent system we named H2A.Z-UAB (Ubiquitylation-dependent Auto-Biotinylation) to directly purify H2A.Zub nucleosomes. H2A.Z-UAB involves expression of a modified form of mammalian H2A.Z that undergoes auto-biotinylation upon ubiquitylation, thereby enabling specific purification of H2A.Zub using streptavidin-coupled reagents. In this study, we validated the effectiveness of this system and demonstrated its versatility in both biochemical assays and genome-wide analyses. Our findings further provide direct biochemical evidence illustrating the preferential association of H3K27me3 and exclusion of H3K4me1/2/3 with ubiquitylated H2A.Z within the nucleosomal context, alongside the preferential association of active chromatin marks with non-ubiquitylated H2A.Z (H2A.Znon-ub). By high-throughput sequencing of the DNA purified from these nucleosomes, which is equivalent to native ChIP-seq, we delineated genomic regions enriched for H2A.Zub or H2A.Znon-ub and corroborated our biochemical findings through genomic co-localization studies. Lastly, our analyses not only showed enrichment of H2A.Zub at transcriptionally inactive genes and silenced chromatin at large but also demonstrated specific enrichment of H2A.Zub at DNA methylated CpG islands and at homopurine-homopyrimidine sequences. Together, our findings illustrate the universal association of H2A.Zub with transcriptionally silenced chromatin across multiple contexts of repression.

## MATERIAL AND METHODS

### Cell culture, transfection, plasmids

HEK293T cells were cultured in Dulbecco’s modified Eagle’s medium (Multicell 319-005-CL) supplemented with 10% fetal bovine serum (Multicell 098150) and penicillin/streptomycin (Multicell 450-201-EL). Transfections were performed using polyethyleneimine (Polysciences 23966). All expression constructs were derived from the pcDNA 3.1 (+) backbone (Invitrogen), with the Flag-BirA fused in-frame to the C-terminus of H2A.Z.1 or H2A.Z.1-K3R3, designated as H2A.Z-FB and H2A.Z-FB-K3R3, respectively. In K3R3 mutant, all three C-terminal ubiquitylation sites on H2A.Z are mutated to arginines (K120/121/125 to R120/121/125). Additionally, the Avi-tag was incorporated in-frame at the N-terminus of ubiquitin, referred to as Avi-ub. A non-biotinylatable Avi-ub was created by converting K10 of the Avi-tag sequence (GLNDIFEAQKIEWHE) to R10. In co-transfection experiments, the plasmid ratio of H2A.Z-FB to Avi-Ub was maintained at 3:1.

### Whole cell and nuclear extractions

Extractions were done as previously described (23). Briefly, cells were harvested by trypsinization, washed in PBS, and pelleted by centrifugation. Cell pellets were then lysed in boiling 2xSDS sample buffer (20mM Tris-HCl pH7.4; 20mM EDTA; 2% SDS; 20% glycerol), and samples were sonicated before storage at −20°C.

### Immunoblotting

All protein samples underwent electrophoresis in 15% SDS-PAGE (4% stacking) in Tris-Glycine-SDS buffer. For Coomassie staining, gels were immersed in staining solution for 30 min to 1 hour, followed by destaining for 1-2 hours, and finally scanned for visualization. For Western blotting, proteins were transferred onto methanol activated PVDF membranes (Millipore IPVH00010) under semi-dry conditions using 1x Towbin buffer via ECl semi-dry transfer unit (Hoefer TE-77X). Membranes were blocked in 1x TBS with 5% milk, except for Avi-HRP, which was blocked in 1x TBS with 5% BSA. Following blocking, membranes were incubated in primary antibody solution (diluted in 1x TBST with 2% milk or BSA; see Table 1 for concentrations) for either 1 hour at room temperature or overnight at 4°C. After primary antibody incubation, membranes were washed three times in 1x TBST for 5 minutes each and then incubated in an appropriate secondary antibody solution (diluted in 2% milk or BSA; 1:10,000) for 1 hour at room temperature. Subsequent to another round of washing as described, blots were soaked in ECL substrate (Millipore WBLUC0500 or WBKLS0500) and developed using a film processor (Konica-Minolta SRX-101A).

### Mononucleosome affinity purification

Generation of mononucleosomes followed a previously established protocol (25). HEK293T cells were cultured in 15 cm-diameter plates and transfected with various constructs as per experimental requirements. After 48 hours post-transfection, cells were washed with 1X PBS and harvested by scraping. All the centrifugation steps were conducted at 4°C. Cells were pelleted at 300 x g for 5 min. Cellular pellets were resuspended in buffer A (20 mM HEPES, pH 7.5, 10 mM KCl, 1.5 mM MgCl_2_, 0.34 M sucrose, 10% glycerol, 1 mM dithiothreitol, 5 mM sodium butyrate, 10 mM NEM, and protease inhibitors), pelleted at 600 x g for 5 min, and then resuspended in buffer A containing 0.2% Triton X-100. The suspension was incubated on ice for 5 minutes. The released nuclei were then centrifuged at 1300 x g for 5 min, washed once in buffer A, and resuspended in cutting buffer (15 mM NaCl, 60 mM KCl, 10 mM Tris pH 7.5, 5 mM sodium butyrate, 10 mM NEM, and protease inhibitors) supplemented with 2 mM CaCl_2_. Micrococcal nuclease (MNase; Worthington 9013-53-0) was added at a concentration of 10 units/1.0 x 10^7^ cells and incubated at 37°C for 30 minutes. The reaction was stopped by adding 20 mM EGTA. The MNase-digested nuclei were centrifuged at 1300 x g for 5 min, and the resulting supernatant (S1) was saved. The digested nuclear pellet was further subjected to hypotonic lysis by resuspension in TE buffer (10 mM Tris-HCl, pH 8.0, 1 mM EDTA) and incubated at 4°C rocking for 1 hour. After centrifugation at 16,000 x g for 5 min, the supernatant (S2) was transferred to a new tube. Salt concentration in S1 and S2 was adjusted to 150 mM NaCl using 2X buffer D and 3X buffer E, respectively. Insoluble material was pelleted via centrifugation, and the clarified supernatants were combined for affinity purification. Streptavidin-agarose (Sigma S1638) or Flag M2-agarose beads (Sigma A2220) were added and incubated overnight at 4°C on an end-over-end rotator. Beads were washed 4 times in 1X Buffer D, followed by 3 washes in 1X Buffer D containing 0.5% Triton X-100. Proteins were eluted from the beads by resuspension in 2X SDS sample buffer and boiled for 10 minutes. For Western blot analysis, samples were run on SDS-polyacrylamide electrophoresis gels following standard procedures.

### ChIP-sequencing

Mononucleosome affinity purification was conducted in duplicate following the protocol described above. Streptavidin-agarose, Flag M2-agarose, or H3 antibody (pulled down using protein G-coupled Dynabeads; Invitrogen) were utilized, and elution was carried out in buffer D containing 1% SDS by end-over-end rotation at room temperature for 2 cycles of 10 minutes each. Following elution, DNA underwent treatment with RNase A and proteinase K, purification via phenol-chloroform extraction, and subsequent re-precipitation with ethanol before being resuspended in water. DNA was then converted to libraries using the Illumina TruSeq ChIP-seq kit and sequenced on an Illumina NextSeq500 in single-end mode.

### ChIP-qPCR

Mononucleosome affinity purification (SA-IP and Flag-IP) was performed from H2A.Z-FB + Avi-Ub transfected cells, following the established protocol. DNA extraction was conducted using the same procedure as for ChIP-seq, and DNA concentrations were determined using Qubit. Ten promoter-overlapping regions from both H2A.Zub and H2A.Znon-ub peak sets, each exhibiting the highest DiffBind fold enrichment, were selected for validation by ChIP-qPCR. For Fig. 3C and Fig. 6C, equal amounts of SA-IP and Flag-IP DNA were analyzed by qPCR. Locus-specific signal was expressed as SA-IP relative to Flag-IP to assess differential representation between the two purified nucleosome pools. For Fig. 4C and Fig. 7C, equal volumes of H3K4me3, H3K27me3 or PRDM1-immunoprecipitated DNA and corresponding input samples were analyzed by qPCR. Enrichment is presented as percent input. Quantitative polymerase chain reactions (qPCR) were arranged in triplicate using either Perfecta SYBR supermix (VWR 95054-02K) or GB Amp Universal SYBR Green qPCR mix (GeneBio Q711-01), accompanied by gene-specific primers. The reactions were run on an Opticon 2 thermocycler (Biorad). Primer sequences can be found in Table 1.

### MeDIP-qPCR

Genomic DNA was extracted from ∼3.2 x 10^6^ untransfected HEK293T cells using an extraction kit (Thermo Scientific K0721). MeDIP was performed as described in Guerrero-Bosagna et., 2015 with the following modifications: (1) sonication of genomic DNA was performed using Fisherbrand™ Model 705 Sonic Dismembrator; (2) anti-methylcytosine antibody from Active Motif (61479) and Protein G-coupled Dynabeads (Invitrogen 10004D) were used; (3) DNA was extracted using phenol-chloroform extraction instead of using spin-filtering columns. Ten regions from both H2A.Zub and H2A.Znon-ub peak sets, each exhibiting the highest DiffBind fold enrichment for which primers could be designed, were selected for amplification. Equal volumes of MeDIP and input DNA, were amplified using primers from either peak set. MeDIP and input samples were run on the same plate for consistency and comparison. Quantitative polymerase chain reactions (qPCR) were arranged in triplicate using GB Amp Universal SYBR Green qPCR mix (GeneBio Q711-01), accompanied by gene-specific primers. The reactions were run on an Opticon 2 thermocycler (Biorad). Enrichment is presented as percent input. Primer sequences can be found in Table 1.

### Computational analysis

#### ChIP-seq analysis

The following steps were performed using Galaxy platform (The Galaxy Community, 2022). Raw FASTQ files were mapped to hg38 human genome assembly using Burrows-Wheeler Aligner (BWA v0.7.17) in Simple Illumina mode. Non-uniquely mapped reads were removed by filtering out reads with mapping quality less than 20 using Filter SAM or BAM (SAMtools v1.9). Read coverage distribution for SA-IP and Flag-IP were prepared in bigWig format using bamCompare or bamCoverage (DeepTools v3.3.2). Peaks were called using MACS2 v2.1.1, wherein band width was set to 150 and upper mfold bound to 100. To identify differentially bound regions enriched in H2A.Zub or H2A.Znon-ub, we performed DiffBind v2.10.0 between SA-IP and Flag-IP, using H3 ChIP-seq as input control (FDR < 0.05).

#### General analysis

Genome browser view of the example peaks was prepared using karyoploteR v1.36.0 (Bioconductor). All the bar charts and boxplots were prepared using ggplot2. Peaks were assigned with annotated genomic features using ChIPseeker v1.30 (R). GO enrichment analysis was performed by ShinyGO v085. CpG rich islands (CGIs) were retrieved from University of California at Santa Cruz (UCSC) and classified as intragenic or intergenic based on −/+100bp overlap with a UCSC annotated gene. Promoter CGIs were defined as overlapping −/+100bp with an annotated TSS. Promoter CGIs that were common with intragenic CGIs were excluded from the latter. Profile plots and heatmaps were generated using plotProfile and plotHeatmap (Deeptools v3.3.2). hPu/hPy-tract coordinates were obtained from the Andre Nussenzweig laboratory based on repeat annotations described in Matos-Rodrigues et al. (2022) (26).

#### Publicly available data

Following publicly available genome-wide sequencing data were used in this study: (a) H3K4me3 ChIP-seq (GSM5395708), H3K4me1 ChIP-seq (GSM5395704), H3K27ac ChIP-seq (GSM5395703) (27); (b) H3K27me3 ChIP-seq (GSM4301076) (28); (c) H3K9me3 ChIP-seq (ENCLB555ACF) (ENCODE Project Consortium); (d) WGBS-seq (GSM1254259) (29); RNA-seq (GSM3734788) (30).

## RESULTS

### Development and characterization of a Ubiquitylation-dependent Auto-Biotinylation (UAB) system for studying monoubiquitylated H2A.Z

We previously developed a method named BICON to specifically purify phosphorylated H3 from total chromatin (31). That method involved fusing the H3 kinase MSK1 to the *E. coli* biotin ligase BirA and leveraged BirA’s specific biotinylation of a 15-amino acid sequence (named Avi-tag) added to H3 to co-modify (phosphorylate and biotinylate) MSK1-targeted nucleosomes. To adapt that approach for studying H2A.Z ubiquitylation, we fused BirA to the C-terminus of H2A.Z and placed the Avi-tag on the N-terminus of the ubiquitin molecule (Avi-ub; Fig. 1A). We also added a Flag-tag peptide sequence between the H2A.Z and BirA domains to facilitate its detection and purification, resulting in an H2A.Z-Flag-BirA fusion, hereafter termed H2A.Z-FB. By co-expressing H2A.Z-FB and Avi-ub in transfected cells, we reasoned that when H2A.Z-FB is monoubiquitylated with Avi-ub, the BirA at its C-terminus would preferentially and automatically biotinylate the Avi-tag on the conjugated ubiquitin since the enzyme and substrate are now physically on the same molecule (Fig. 1A). Therefore, we essentially generated a version of H2A.Z that self-biotinylates when it is ubiquitylated with Avi-ub.

**Figure 1.**
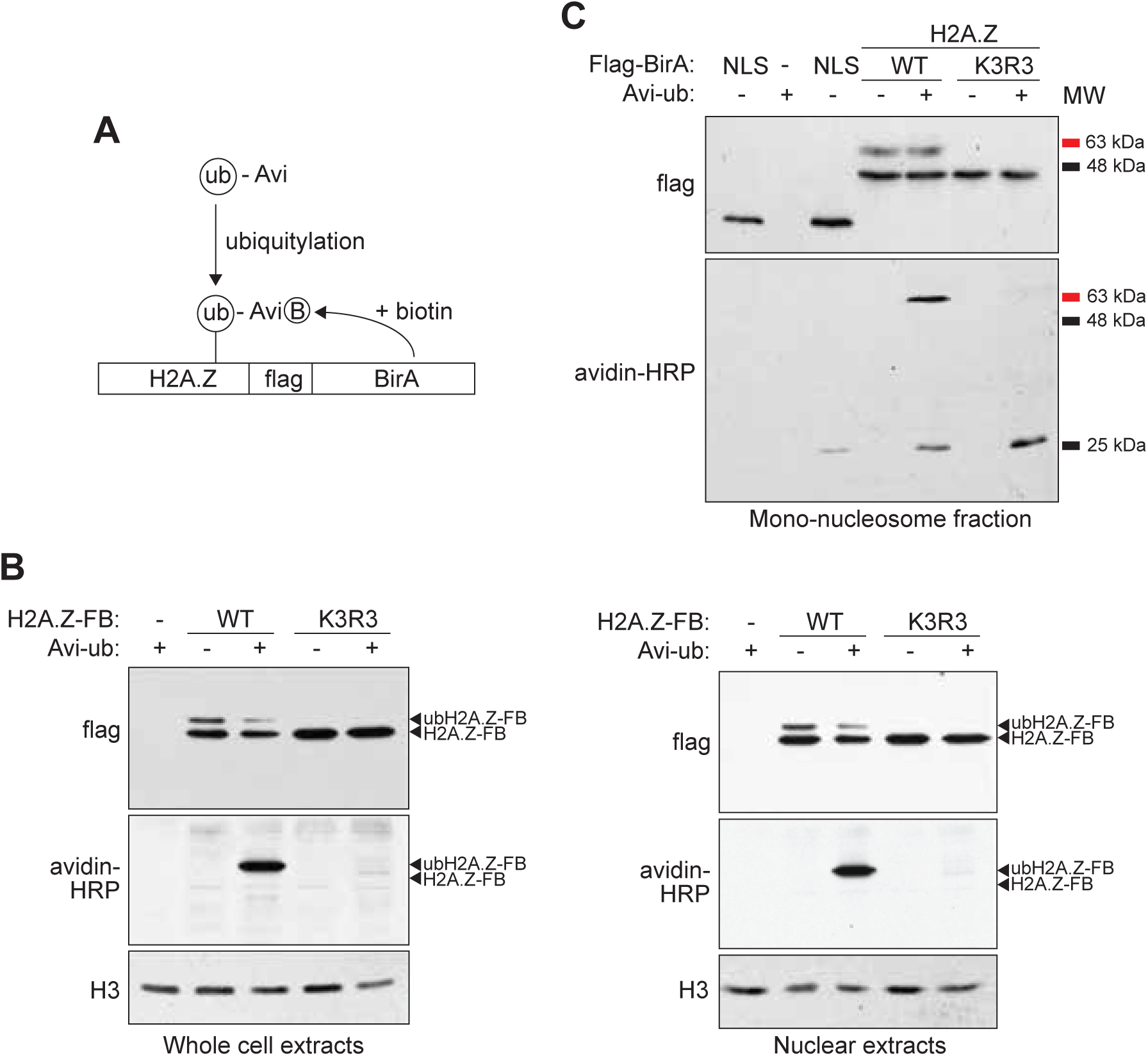
Validation of the auto-biotinylation of Avi-ubiquitylated H2A.Z-Flag-BirA. **A.** Schematic representation of the H2A.Z-Flag-BirA (H2A.Z-FB) fusion and its auto-biotinylation upon ubiquitylation with Avi-tagged ubiquitin (Avi-Ub). **B.** HEK293T cells were co-transfected with wild-type (WT) or non-ubiquitylatable (K3R3) H2A.Z-FB with or without Avi-Ub. Whole-cell or nuclear extracts were harvested from the transfected cells, and total H2A.Z-FB or biotinylated H2A.Z-FB were detected by Western blots using Flag antibody or horse-radish peroxidase-coupled avidin (Avidin-HRP). H3 served as a loading control. **C.** Mononucleosomes were prepared from HEK293T cells transfected with the indicated NLS-Flag-BirA, H2A.Z-FB (WT or K3R3) and Avi-Ub combinations. The presence of Flag tagged and biotinylated proteins were detected by Western blots using Flag antibody or Avidin-HRP. The molecular weight (MW) markers show the relative sizes of the bands detected by the Western blots.

As a proof of concept, we first co-transfected H2A.Z-FB and Avi-ub expression constructs into human HEK293T cells and examined their expression and biotinylation by Western blot analysis (Fig. 1B). As an additional control, we generated a non-ubiquitylatable mutant form of H2A.Z-FB in which the known sites of H2A.Z monoubiquitylation (K120, K121, and K125) are mutated to arginines (referred to as the K3R3 mutant) and co-expressed it with Avi-ub in HEK293T cells. Western blot of whole cell extracts or nuclear extracts of the respective transfected cells using anti-Flag antibody showed strong expression of both WT- and K3R3-H2A.Z-FB (Fig. 1B). As expected, only the WT-H2A.Z-FB sample contained a shifted band corresponding to monoubiquitylated H2A.Z-FB. We next examined the presence of biotinylated proteins in the same extracts using horse-radish peroxidase-conjugated avidin (Avidin-HRP) and found that the predominant and most abundant biotinylated protein was indeed the Avi-ub-H2A.Z-FB, which co-migrated with the shifted higher molecular weight band detected in the Flag Western blots (Fig. 1B). The Avidin-HRP-detected band in the WT-H2A.Z-FB co-transfection was not present in the K3R3-H2A.Z-FB mutant co-transfection, further confirming that biotinylation is specific only to the Avi-ubiquitylated H2A.Z-FB. These results indicate that our experimental system works as predicted and that Avi-ub-H2A.Z-FB is the predominant biotinylated protein in the transfected cells.

Further characterization of this system showed that within mononucleosome-enriched fractions (harvested from micrococcal-nuclease digested nuclei), in addition to Avi-ub-H2A.Z-FB, Avidin-HRP also detected a small amount of a lower molecular weight band (∼ 25 kDa), which likely corresponds to endogenous ubiquitylated H2A.Z (Fig. 1C). Since H2A.Z-FB can theoretically partner with endogenous H2A.Z (or H2A) in the nucleosomal context, it is possible that it also biotinylates nearby Avi-ub-H2A.Z/H2A, either on the same nucleosome or in close proximity. Consistent with this, the lower band was also observed in cells transfected with the NLS-Flag-BirA control (BirA fused to Flag and a nuclear localization signal), indicating that co-expression of BirA and Avi-ub alone can lead to low levels of background biotinylation of Avi-ubiquitylated endogenous histones. Nevertheless, this background amount is minimal compared to the strong biotinylation of Avi-ub-H2A.Z-FB in cells co-transfected with H2A.Z-FB and Avi-ub. Notably, in cells expressing the K3R3-H2A.Z-FB mutant along with Avi-ub, only the lower band is detected by Avidin-HRP, and it was more prominent than in the wild-type condition. Overall, these findings demonstrate wild-type H2A.Z-FB preferentially biotinylates the Avi-tagged ubiquitin on itself, but in the absence of an intra-molecular target site, BirA can shift to biotinylate other nearby Avi-tagged substrates.

### Affinity purification of mononucleosomes containing Avi-ubiquitylated H2A.Z-FB

Previous work by us and others already showed that epitope- or GFP-tagging of H2A.Z at the C-terminus does not interfere with its incorporation into nucleosomes, so we anticipated similar behaviour with the inclusion of the BirA tag (23,32). To test this, we harvested micrococcal nuclease-digested chromatin from cells co-transfected with H2A.Z-FB and Avi-ub and isolated mononucleosomes by affinity purification using streptavidin (SA) or anti-Flag antibody-coupled beads (hereafter referred to as SA-IP and Flag-IP, respectively) (Fig. 2A). As shown in the Coomassie-stained gel (Fig. 2B), both SA- or Flag-IP samples contained stoichiometric amounts of core histones (i.e., roughly equal amounts of H2B, H3 and H4) that co-purified with H2A.Z-FB or H2A.Z-K3R3 (lanes 6, 7 and 8 of Fig. 2B). This confirms that both non-ubiquitylated and ubiquitylated H2A.Z-FB, and the mutant H2A.Z-K3R3 are incorporated into nucleosomes, and that intact nucleosomes are efficiently purified by the SA- or Flag-IP. As an additional control, we co-expressed H2A.Z-FB with a K10R mutant form of Avi-ub whereby the lysine residue within the Avi-tag (required for biotinylation) was mutated to arginine. When mononucleosomes harvested from these cells were subjected to SA-IP, no detectable core histones were co-purified. This result verifies that core histone co-purification with Avi-ub-H2A.Z-FB is due to its incorporation into nucleosomes, and not due to non-specific binding to SA-beads.

**Figure 2.**
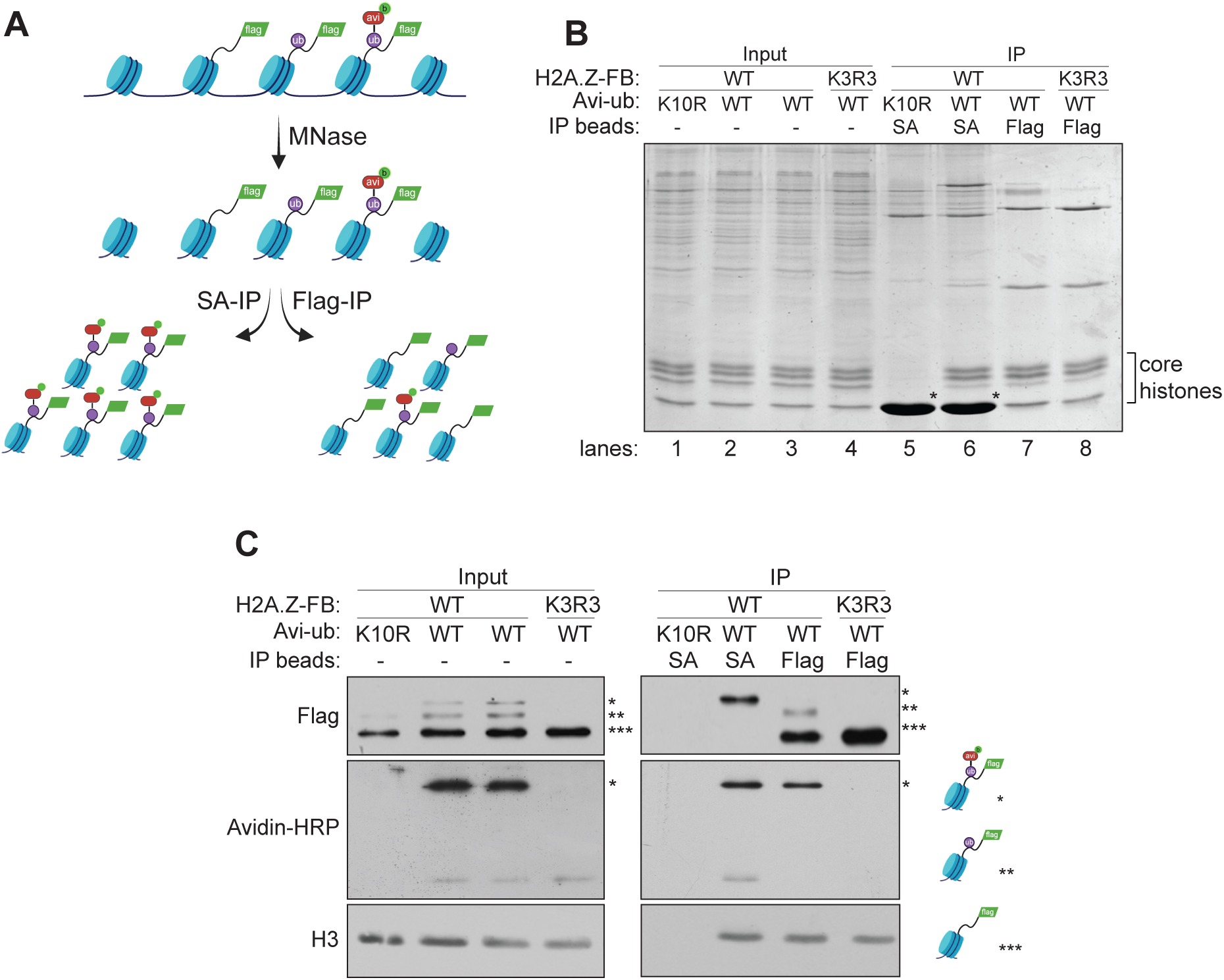
Separation of distinct pools of H2A.Z-FB nucleosomes by streptavidin (SA-IP) or Flag antibody-based (Flag-IP) affinity purification. **A.** Schematic diagram illustrating the expected types of H2A.Z-FB nucleosomes present in H2A.Z-FB + Avi-Ub transfected cells and the distinct nucleosome populations isolated by SA- or Flag-IP. **B.** Coomassie-stained gel of the input mononucleosome fraction and SA- or Flag-purified nucleosomes. * indicates non-specific bands often seen in SA-IPs. **C.** Western blot analysis of the input and affinity-purified mononucleosomes using Flag antibody and Avidin-HRP. The different types of ubiquitylated and non-ubiquitylated H2A.Z-FB (illustrated on the right-hand side and marked by single, double, and triple asterisks) were resolved on long SDS polyacrylamide gels and detected by Flag antibody.

We next used Western blot analyses to characterize the SA- or Flag-purified mononucleosomes from cells expressing various combinations of H2A.Z-FB and Avi-ub (Fig. 2C). By using higher percentage polyacrylamide gels with extended run times to better resolve H2A.Z-FB species (compared to those shown in Fig. 1), we were able to separate and detect non-ubiquitylated H2A.Z-FB (the fastest migrating band), H2A.Z-FB ubiquitylated with endogenous ubiquitin (the middle band), and those ubiquitylated with Avi-ub (the slowest migrating top band; Fig. 2C). Crucially, when we normalized the purified nucleosomes based on their H3 levels, we observe that SA-IP specifically pulled down the Avi-ub-H2A.Z-FB, whereas Flag-IP primarily pulled down the non-ubiquitylated H2A.Z-FB and a smaller amount of H2A.Z-FB monoubiquitylated with endogenous (non-Avi-tagged) ubiquitin. As expected, in cells expressing the H2A.Z-FB K3R3 mutant and Avi-ub, the Flag-IP pulled down only non-ubiquitylated H2A.Z-FB. Similarly, in cells expressing WT H2A.Z-FB along with the K10R Avi-ub mutant, the SA-IP did not pull down any H2A.Z-FB or endogenous histones, further confirming the specificity of SA-based affinity purification. Finally, we note that SA-IP predominantly pulled down the Avi-ub-H2A.Z, with only a minor amount of Avi-ubiquitylated endogenous H2A.Z/H2A also detected (Fig. 2C, right panel). Taken together, these results demonstrate that our experimental system, combined with SA-based purification strategy, provide robust enrichment of ubiquitylated H2A.Z-containing nucleosomes, thereby enabling more in-depth studies of their biochemical and genomic characteristics.

### Genomic distribution profile of ubiquitylated and non-ubiquitylated H2A.Z

Having confirmed successful purification of Avi-ub-H2A.Z-FB (hereafter referred as H2A.Zub) containing nucleosomes, we proceeded to perform ChIP-seq on the nucleosomal DNA to map the genomic distribution of this mark. Using conditions equivalent to ChIP, we extracted DNA from the SA- or Flag-purified nucleosomes (from biological replicate preparations) and performed high-throughput sequencing. In parallel, we also performed native ChIP-seq using an antibody against H3 as a control for histone density. Overall, we identified approximately 340,000-360,000 peaks in both SA- and Flag-IP samples, with a substantial (∼70%) overlap between them. This degree of overlap is expected, as the Flag-IP captures both non-ubiquitylated and ubiquitylated H2A.Z-FB, whereas the SA-IP specifically enriches for Avi-ub-H2A.Z-FB. We note that the SA-IP peaks are not entirely contained within the Flag-IP peak set, which may be due to the low abundance of H2A.Zub in cells and the higher affinity of streptavidin compared to the Flag antibody. As a result, streptavidin beads may be more effective at capturing low-abundance H2A.Zub nucleosomes, which could otherwise be outcompeted by more abundant non-ubiquitylated nucleosomes in the Flag-IP samples.

H2A.Z has been reported to be enriched at the +1, and to a lesser extent at the −1 nucleosomes flanking transcription start sites (TSSs) of many genes (17). When the SA- and Flag-IP signals were mapped to TSSs, we indeed observed a strong peak at +1 nucleosome position and a weaker peak at the −1 nucleosome position for both IP types (Fig. 3A, top panels), indicating that H2A.Z-FB is correctly targeted to the endogenous H2A.Z sites within the genome. Interestingly, when we mapped SA- and Flag-IP signals to genes stratified by expression levels (no, low, or high expression based on RNA-seq from 293T cells), SA-IP enrichment at the TSSs was comparable across all groups. In contrast, Flag-IP signal was significantly higher at the TSSs of both low- or high-expression genes compared to non-expressed genes (Fig. 3A, middle panels). This difference became more apparent when the SA-IP signal was normalized to the Flag-IP signal (Fig. 3A, bottom panel), revealing a relative depletion of SA-IP signal only at low- and high-expression genes. These findings suggest that while similar amounts of Avi-ub-H2A.Z-FB are present at the +1 nucleosome irrespective of gene expression status, total H2A.Z-FB (predominantly non-ubiquitylated) is significantly more enriched at the +1 and −1 nucleosome positions of active genes. Notably, Flag-IP signal levels flanking TSSs are nearly identical between low- and high-expressing genes, suggesting that while active promoters generally contain more H2A.Z than inactive promoters, the absolute H2A.Z levels at +1/−1 nucleosomes do not correlate with expression magnitude (i.e., low vs. high expression).

**Figure 3.**
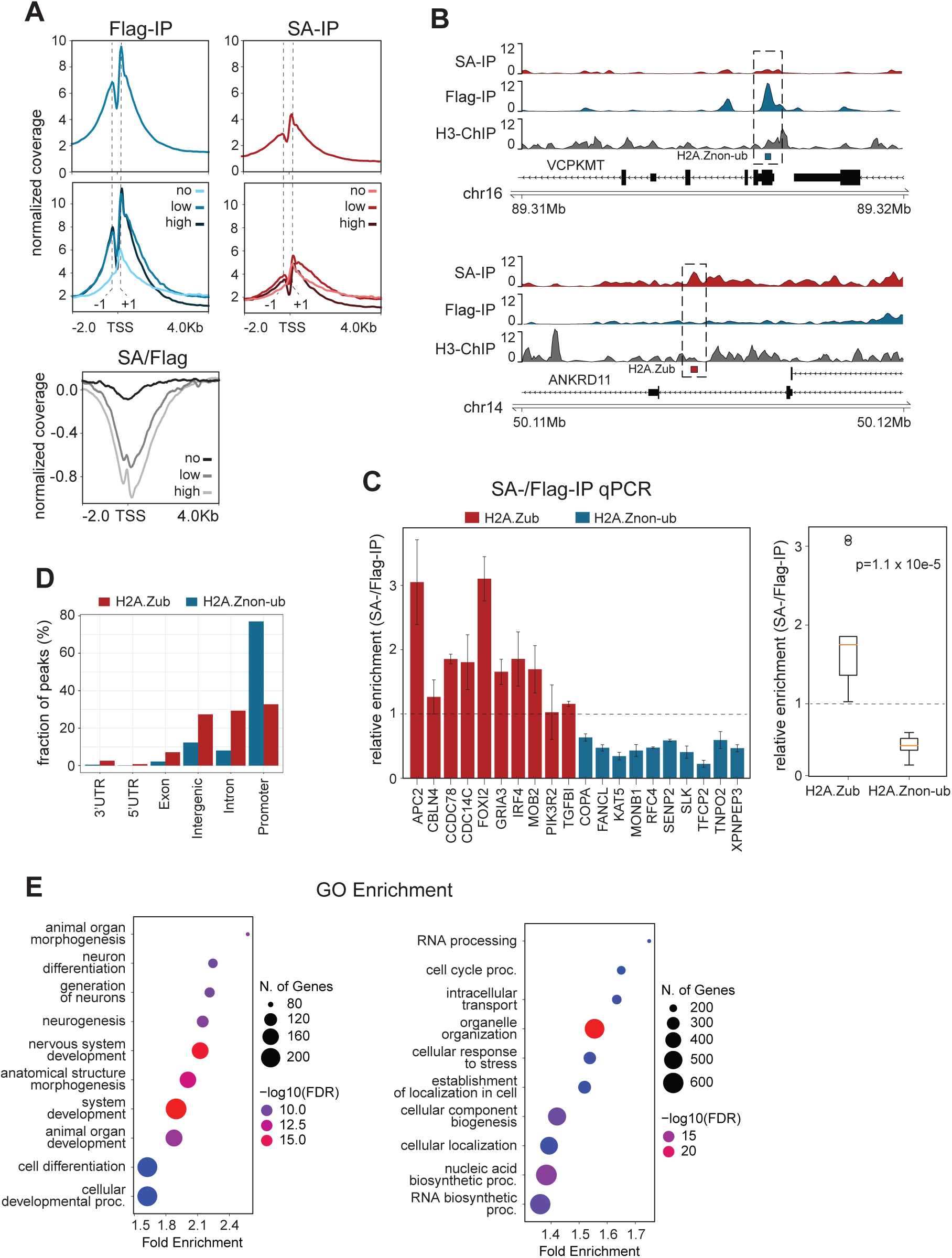
Genomic analysis of ubiquitylated and non-ubiquitylated H2A.Z containing nucleosomes. **A.** Profile plots representing Flag-, SA- or SA/Flag-IP signals over all TSSs. For middle and bottom panels TSSs are classified based on RNA-Seq TPM values; high expression (>10), low expression (1 to 10), no expression (<1). **B.** Genomic browser view of example peaks from each enriched peak set. **C.** Validation and comparison of SA-IP normalized over Flag-IP signal at identified H2A.Zub and H2A.Znon-ub peaks by qPCR (Mann-Whitney U test). Ten peaks from the respective sets were chosen based on an overlap with UCSC annotated promoters. Relative enrichment values >1 indicate higher SA-IP signal, whereas values <1 indicate higher Flag-IP signal. **D.** Genome-wide distribution of identified peaks over UCSC annotated genomic features. **E.** ShinyGO enrichment analysis illustrating significant biological processes associated with each peak set (FDR q<0.05; significant by both binomial and hypergeometric tests).

To identify regions specifically enriched for either ubiquitylated or non-ubiquitylated H2A.Z, we utilized DiffBind (Bioconductor) to conduct differential enrichment analysis between SA-and Flag-IP peaks, using H3 ChIP as an input control. This analysis revealed 3,197 peaks differentially enriched in SA-IP nucleosomes and 3,561 peaks enriched in Flag-IP nucleosomes. We classified the former as H2A.Zub-specific peaks and the latter as H2A.Znon-ub-specific peaks. Genome browser visualization confirmed the differential enrichment of SA-IP versus Flag-IP ChIP-seq signals for representative peaks from each set (Fig. 3B). To validate the ChIP-seq results, we performed gene-specific SA- and Flag-IP qPCR on ten representative promoters from each group and compared their amplification levels (Fig 3C). For each promoter, we calculated the relative enrichment of the SA-IP qPCR signal by normalizing it against Flag-IP qPCR signal. As shown in the combined analysis, the normalized SA-IP values were greater than 1 at H2A.Zub peaks and less than 1 at H2A.Znon-ub peaks, thereby confirming the differential enrichment observed in the ChIP-seq data (Fig. 3C). Next, we investigated the genomic distribution of these enriched peaks across annotation classes (Fig. 3D). The majority of the H2A.Znon-ub peaks were found on promoters, consistent with the known distribution of total H2A.Z in the genome (17). In contrast, H2A.Zub peaks displayed a more even distribution across promoters, introns, and intergenic regions.

To explore whether genes associated with H2A.Zub and H2A.Znon-ub are functionally distinct, we carried out Gene Ontology (GO) enrichment analysis on the promoter-associated peaks in each group. H2A.Znon-ub genes were enriched for core biological functions, such as metabolism, nucleic-acid processing, cell-cycle regulation, transcription, and translation (Fig. 3E). In contrast, H2A.Zub genes were predominantly enriched for processes related to cellular or organismal development and differentiation (Fig. 3E). These findings align with the well-established role of H2A.Z in active transcription, as well as the enrichment of H2A.Zub at bivalent promoters of developmental regulators in mouse embryonic stem cells (mESCs) (3,6,21,24). In summary, the integration of ChIP-seq with our ubiquitylation-dependent self-biotinylating system enabled identification of genomic regions enriched for either ubiquitylated or non-ubiquitylated H2A.Z, facilitating a comparative analysis based on their modification status.

### Ubiquitylated H2A.Z associates with repressive chromatin states

Having established that distinct pools of H2A.Z-FB nucleosomes are captured by the SA-and Flag-IPs, we next examined the histone modification patterns on these nucleosomes by Western blot. Tested histone marks include those that commonly define active vs. silenced chromatin. As before, we normalized the samples based on their H3 levels and confirmed that SA-IP preferentially pulls down H2A.Zub nucleosomes, while Flag-IP predominantly captured non-ubiquitylated H2A.Z nucleosomes (Fig. 4A). Comparison of the two nucleosome pools revealed similar overall levels of H3K9me2 and H3K9me3 (repressive chromatin marks associated with heterochromatin), as well as H4K16ac (an activating histone acetylation mark) (33–35). However, in contrast to total and non-ubiquitylated H2A.Z nucleosomes, H2A.Zub nucleosomes were notably depleted of H3K4 methylation (particularly H3K4me2 and H3K4me3), H3K27ac, and hyperacetylated H4, all which are characteristic marks of transcriptionally active chromatin (36–38). Instead, the H2A.Zub nucleosomes were enriched for H3K27me3, a hallmark of polycomb-silenced facultative heterochromatin (39).

**Figure 4.**
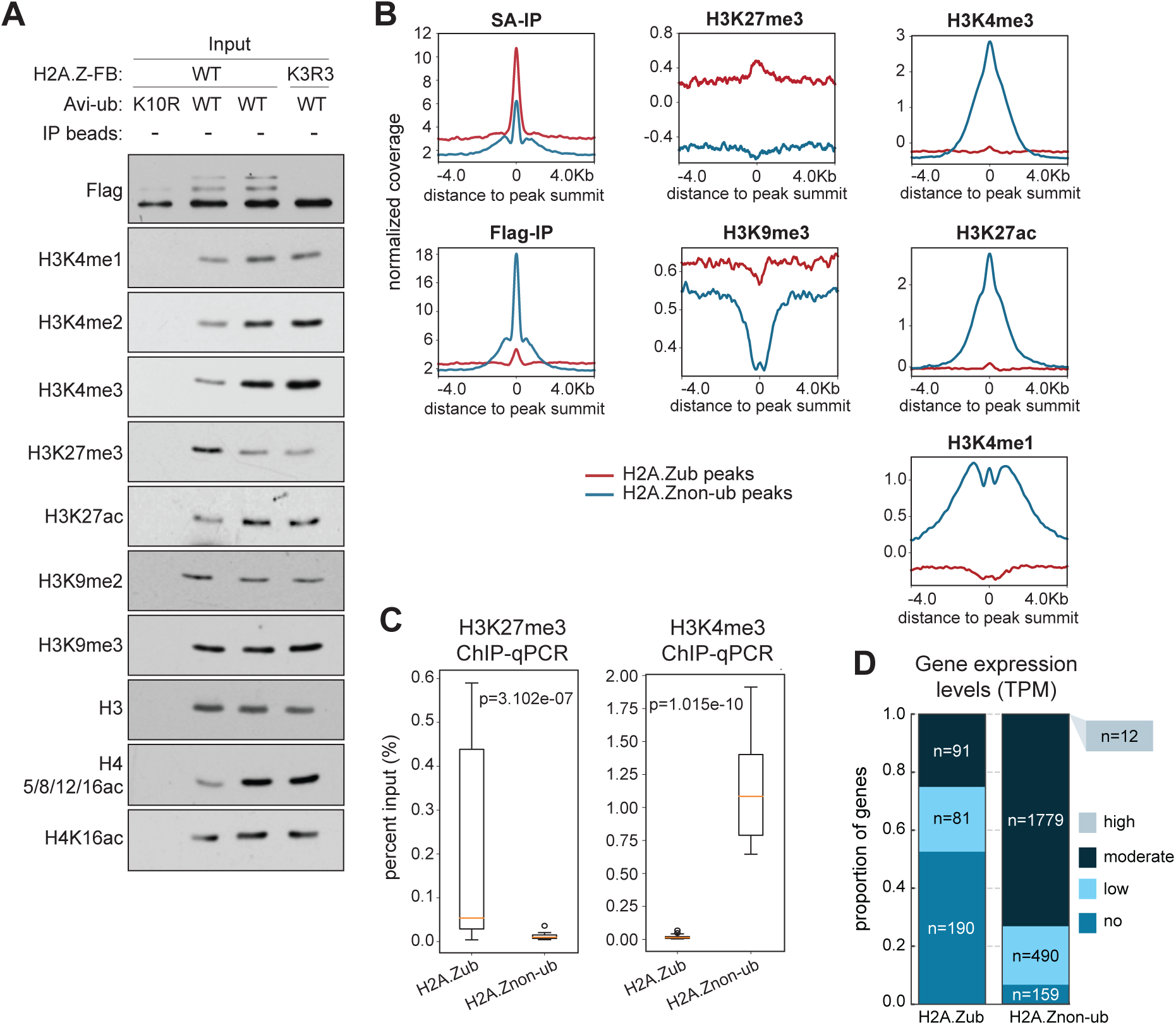
Ubiquitylated H2A.Z associates with repressive chromatin states. **A.** SA- and Flag-IP were performed as described above, and the levels of histone modifications in the input and the captured mononucleosomes were examined by Western blot. H3 serves as a control for chromatin density. **B.** Profile plots showing various ChIP-seq signals over H2A.Zub and H2A.Znon-ub peaks. The signals are centred at the peak summits of either peak set (H2A.Zub peaks, n = 3197; H2A.Znon-ub peaks, n = 3561). **C.** H3K4me3 and H3K27me3 ChIP-qPCR validation and comparison at H2A.Zub (n=10) and H2A.Znon-ub (n=10) associated promoters (Mann-Whitney U test). **D.** Expression levels of genes with promoter-associated H2A.Zub or H2A.Znon-ub peaks were stratified based on RNA-seq TPM values; high expression (>1000), moderate expression (10 to 1000), low expression (1 to 10), no expression (<1).

To further validate these biochemical characteristics, we acquired publicly available HEK293T ChIP-seq data for several of the histone marks studied above and conducted a comparative analysis of their enrichment at H2A.Zub versus H2A.Znon-ub peaks (Fig. 4B). Consistent with the biochemical profiles of H2A.Z nucleosomes, active histone marks such as H3K4me1, H3K4me3, and H3K27ac are highly enriched at H2A.Znon-ub peaks but largely depleted at H2A.Zub peaks. Conversely, the H2A.Zub peaks were located within broad H3K27me3 domains, with the peak of H3K27me3 aligning with the center of H2A.Zub enrichment. In contrast, H2A.Znon-ub peaks showed an inverse correlation with H3K27me3 levels. Lastly, H3K9me3 was depleted at both peak groups, albeit to a greater extent at H2A.Znon-ub peaks. In summary, these genome-wide findings strongly align with the biochemical characteristics of purified H2A.Zub nucleosomes and collectively reveal a general association between H2A.Zub and transcriptionally silent histone PTMs within the nucleosome and chromatin context.

To further explore the transcriptional activity associated with H2A.Zub-linked genes, we compared the expression levels of H2A.Zub and H2A.Znon-ub promoter-associated genes using publicly available RNA-seq data from HEK293T cells (30). When we stratified promoters by expression levels, we found that over 50% of the H2A.Zub promoters showed no expression (defined as <1 TPM), whereas less than 5% of H2A.Znon-ub promoters showed complete absence of expression (Fig. 4D). Conversely, less than 25% of H2A.Zub promoters exhibited moderate or high expression levels (TPM, 10 to 1000 and >1000, respectively), while this number was more than 70% for H2A.Znon-ub promoters. Together, these findings demonstrate a preferential enrichment of ubiquitylated H2A.Z at transcriptionally inactive promoters, and of non-ubiquitylated H2A.Z at active promoters, consistent with their distinct chromatin signatures and functional implications (Fig. 4A-B; Fig. 3E).

### H2A.Zub is enriched at DNA methylated CGIs

While the antagonistic relationship between H2A.Z and DNA methylation is well established, it is unknown whether this anti-correlation extends to ubiquitylated H2A.Z (40,41). To that end, we examined publicly available HEK293 whole-genome bisulfite sequencing (WGBS-seq) data at H2A.Zub and H2A.Znon-ub peaks (Fig. 5A). As expected, H2A.Znon-ub peaks showed marked depletion of WGBS-seq signal, consistent with the known anti-correlation between DNA methylation and total H2A.Z occupancy. Conversely, and notably, there was a pronounced enrichment of DNA methylation at H2A.Zub peaks (Fig. 5A). This positive correlation between H2A.Zub and DNA methylation aligns with previous reports linking DNA methylation to other ubiquitylated histones (42–44).

**Figure 5.**
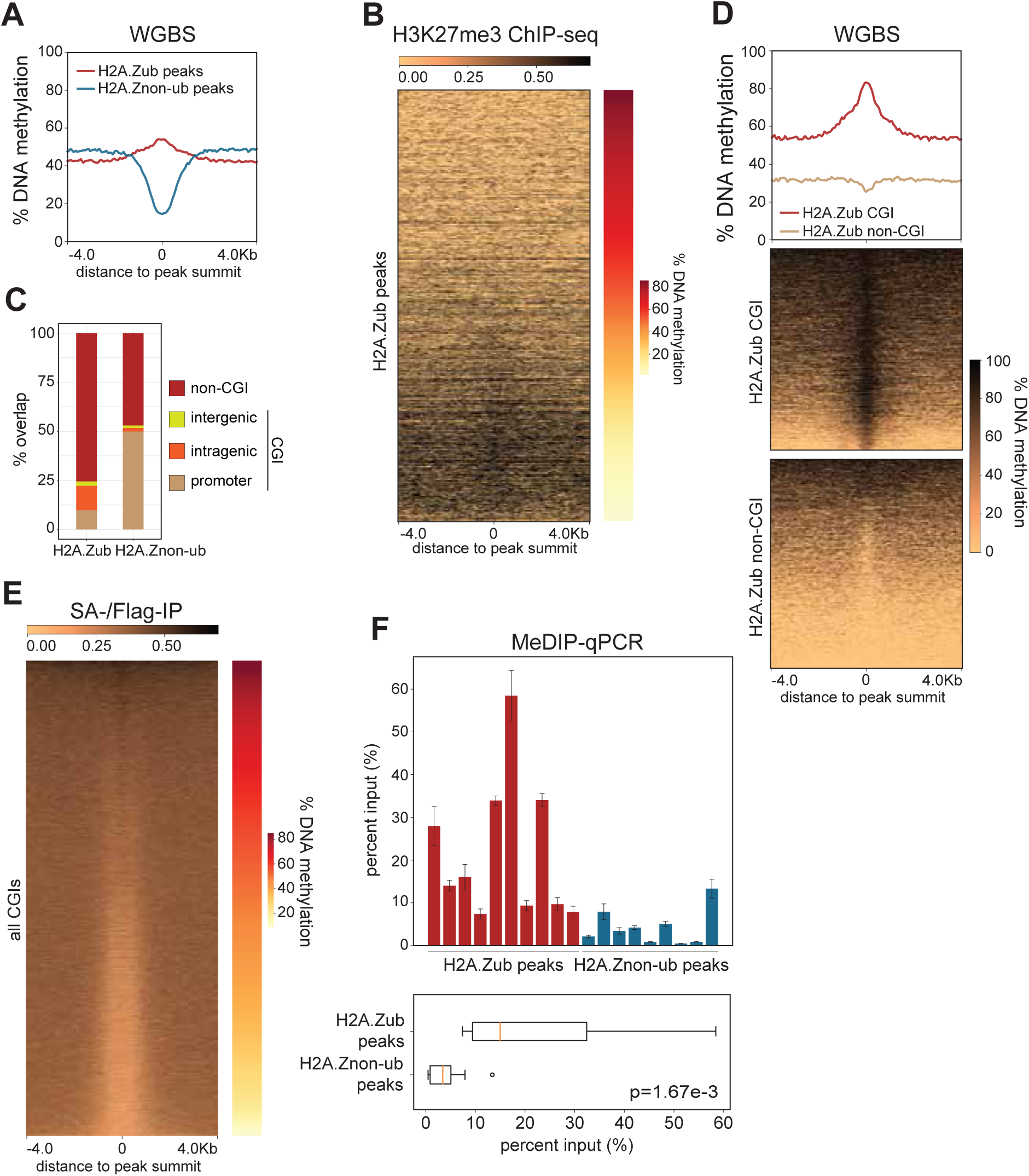
H2A.Zub is enriched at DNA methylated CpG islands (CGIs). **A.** Average CpG methylation profiles generated from WGBS-seq over H2A.Zub and H2A.Znon-ub peaks, centred at their respective peak summits. **B.** Density profile for H3K27me3 at H2A.Zub peaks ranked based on DNA methylation levels. **C.** Fraction of H2A.Zub and H2A.Znon-ub peaks overlapping CpG islands (CGI) as defined by UCSC. **D.** Average CpG methylation profiles over CGI and non-CGI H2A.Zub peaks, centered at their peak summits. **E.** Density profile for SA-/Flag-IP signal representing H2A.Zub levels over all CGIs (n=32,038) ranked based on DNA methylation levels. **F.** MeDIP-qPCR signal is compared between select H2A.Zub (n=10) and H2A.Znon-ub (n=9) peaks (Mann-Whitney U test).

Given the overlap between H3K27me3 and H2A.Zub (Fig. 4B), we sought to investigate the interplay between DNA methylation and H3K27me3 at H2A.Zub peaks. Previous studies have highlighted the mutually exclusive relationship between H3K27me3 and DNA methylation, attributed in part to the preferential deposition of H3K27me3 at CpG islands (CGIs) and the exclusion of DNA methylation from these regions (45,46). To explore this relationship, we ranked H2A.Zub peaks from high to low DNA methylation enrichment based on WGBS-seq signal and examined the corresponding distribution of H3K27me3 ChIP-seq signals (Fig. 5B). H2A.Zub peaks with the lowest DNA methylation enrichment exhibited the highest H3K27me3 signal. Thus, the antagonism between H3K27me3 and DNA methylation persists at H2A.Zub-enriched regions, functionally dividing them into subsets marked predominantly by either modification.

CpG islands (CGIs) are genomic regions with a high density of CpG dinucleotides, typically resistant to DNA methylation and are found at approximately 70% of all vertebrate promoters (47). The divergence in DNA methylation levels between H2A.Zub and H2A.Znon-ub peaks may stem, at least in part, from the higher prevalence of H2A.Znon-ub at promoters relative to H2A.Zub (Fig. 3D). To examine DNA methylation levels in the context of CGIs, we classified H2A.Zub and H2A.Znon-ub peaks based on their overlap with promoter, intragenic or intergenic CGIs as defined by UCSC (Fig. 5C). As anticipated, a larger proportion of H2A.Znon-ub peaks (∼53%) overlapped with CGIs compared to H2A.Zub peaks (∼25%). However, closer inspection of WGBS-seq signal at H2A.Zub peaks revealed that DNA methylation was concentrated almost exclusively within the H2A.Zub CGI regions, whereas non-CGI sites showed minimal signal (Fig. 5D). Notably, the heatmap of H2A.Zub CGI regions showed nearly complete methylation across this group (Fig. 5D, middle panel). Methylation at CpG islands is rare, occurring only in select regions of the genome (48). The observation that many H2A.Zub CGIs (n=∼800) exhibit elevated levels of methylation, whereas H2A.Znon-ub CGIs lack methylation entirely, suggests a potential role for H2A.Zub in the promotion or maintenance of CGI methylation. To further corroborate this association, we ranked all genomic CGIs (n = 32,038) based on their DNA methylation levels and plotted SA-IP/Flag-IP ChIP-seq signals in the same sorted order (Fig. 5E). Strikingly, highly methylated CGIs exhibited elevated SA-IP/Flag-IP signal (the higher portion of the heat map shown in Fig. 5E), indicating that increased H2A.Zub occupancy is a common feature of these regions.

To validate the elevated DNA methylation levels at H2A.Zub sites, we performed methylated DNA immunoprecipitation (MeDIP) in HEK293T cells and assessed DNA methylation levels at H2A.Zub versus H2A.Znon-ub peaks by qPCR (Fig. 5F). As an unbiased approach, regions for qPCR were selected based on highest differential enrichment for either H2A.Zub or H2A.Znon-ub in the original ChIP-seq analysis, independent of their WGBS-seq signal levels. Although not all H2A.Zub peaks examined showed elevated DNA methylation, average enrichment across the group was significantly higher than that of H2A.Znon-ub peaks (Fig. 5F, p=1.67e-3), consistent with the trends observed in the WGBS-seq analysis (Fig. 5A).

### H2A.Zub marks homopurine-homopyrimidine (hPu/hPy) tracts

Upon closer examination of the sequence composition within H2A.Zub peaks, we identified a significant predominance of long uninterrupted runs of homopurine-homopyrimidine sequences (>50 bp) (hPu/hPy-tracts), such as the representative example shown in Fig. 6A.

**Figure 6.**
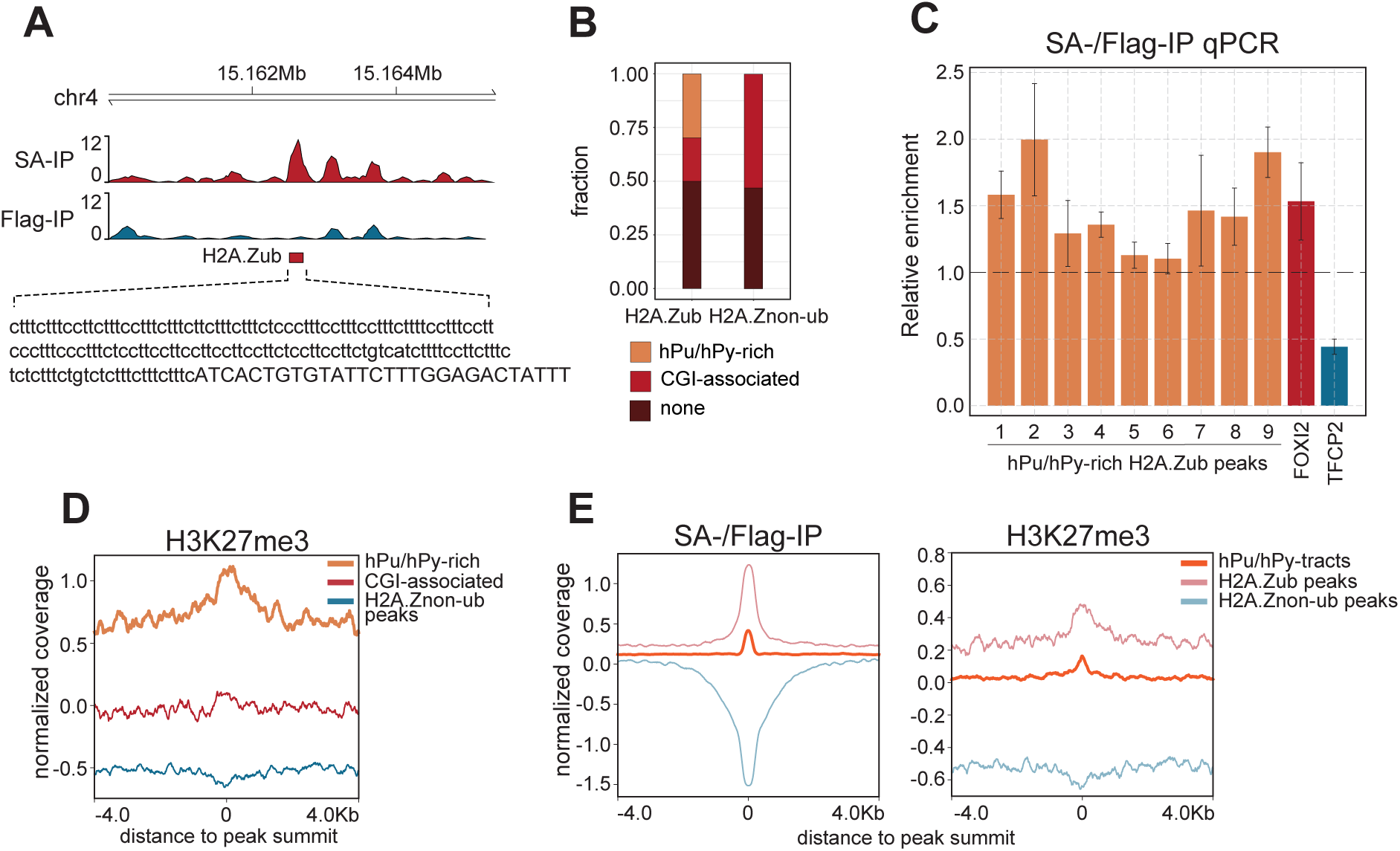
H2A.Zub marks homopurine-homopyrimidine (hPu/hPy) tracts. **A.** Genome browser view of an example hPu/hPy-rich H2A.Zub peak sequence. **B.** H2A.Zub and H2A.Znon-ub peaks are classified based on overlap with CGIs or hPu/hPy-tracts. C. ChIP-qPCR validation of H2A.Zub levels (SA-/Flag-IP) at selected hPu/hPy-rich H2A.Zub peaks. Positive (FOXI2) and negative (TFCP2) controls are validated H2A.Zub and H2A.Znon-ub associated promoters, respectively. **D.** Density profile for H3K27me3 at hPu/hPy-rich or CGI-associated H2A.Zub peaks and H2A.Znon-ub peaks. **E.** Density profiles of H3K27me3 and H2A.Zub (SA-/Flag-IP) over genome-wide hPu/hPy-tracts. Signals over H2A.Zub and H2A.Znon-ub peaks were included for comparison.

Notably, these hPu/hPy-tracts did not correspond to any specific repetitive oligonucleotide motif or defined repeat family (data not shown). To estimate the prevalence of H2A.Zub peaks harboring hPu/hPy-tracts (hPu/hPy-rich H2A.Zub peaks), we selected peaks with purine or pyrimidine content exceeding 75% on either strand and manually confirmed the presence of hPu/hPy-tracts. Using this conservative criterion, we found that more than one-third of H2A.Zub peaks harbored hPu/hPy-tracts, whereas such sequences were almost entirely absent from H2A.Znon-ub peaks (Fig. 6B). Consistent with their distinct sequence compositions, hPu/hPy-rich H2A.Zub peaks rarely overlapped with CpG islands. Accordingly, H2A.Zub peaks can be grouped into three categories: hPu/hPy-rich, CGI-associated, or regions enriched for neither sequence class. We validated H2A.Zub enrichment at selected hPu/hPy-rich peaks by performing SA-/Flag-IP qPCR (SA-IP qPCR normalized to Flag-IP qPCR) (Fig. 6C).

As mentioned previously, DNA methylation and H3K27me3 mark distinct subsets of H2A.Zub peaks (Fig. 5B). Given that DNA methylation is uniquely enriched at CGI-associated H2A.Zub peaks, which do not overlap with hPu/hPy-rich peaks, we examined H3K27me3 enrichment at the latter set of peaks. Indeed, we detected significantly elevated levels of H3K27me3 at hPu/hPy-rich peaks, indicating that these sequences are concomitantly enriched for H2A.Zub and H3K27me3 (Fig. 6D). We then asked whether H2A.Zub and H3K27me3 enrichment is a universal property of hPu/hPy tracts. To this end, we mapped H2A.Zub and H3K27me3 ChIP-seq signals across ∼40,000 hPu/hPy tract coordinates and observed enrichment of both marks relative to H2A.Znon-ub peaks used as a control (Fig. 6E). Given that H2A.Zub and H3K27me3 are Polycomb-associated marks linked to heterochromatin, our results suggest a potential role for heterochromatin formation at hPu/hPy tracts (23,39).

### PRDM1 and H2A.Zub interaction at homopurine-homopyrimidine (hPu/hPy) tracts

H2A.Zub enrichment at a large number of hPu/hPy tracts with a unique sequence composition implicates a potential transcription factor (TF)-mediated, sequence-specific recruitment mechanism. To investigate this possibility, we performed motif enrichment analysis at these sites using SEA from the MEME Suite and found that the majority of the identified TFs have been implicated in transcriptional repression (Fig. 7A). Notably, one of the top hits, PRDM1, has been shown to play a role in the recruitment of PRC1/2 complexes to establish bivalent enhancers during endoderm differentiation (49). Using a publicly available PRDM1 ChIP-seq dataset in HEK293 cells, we evaluated PRDM1 binding at H2A.Zub peaks and observed strong enrichment specifically at hPu/hPy-rich peaks, but not at CGI-associated peaks (Fig. 7B). We also performed PRDM1 ChIP-qPCR in 293T cells and further confirmed enrichment of PRDM1 at hPu/hPy-containing H2A.Zub peaks but not at CGI-associated H2A.Zub peak (FOXI2), thus validating the colocalization of PRDM1 and H2A.Zub at these unique sequences (Fig. 7C).

**Figure 7.**
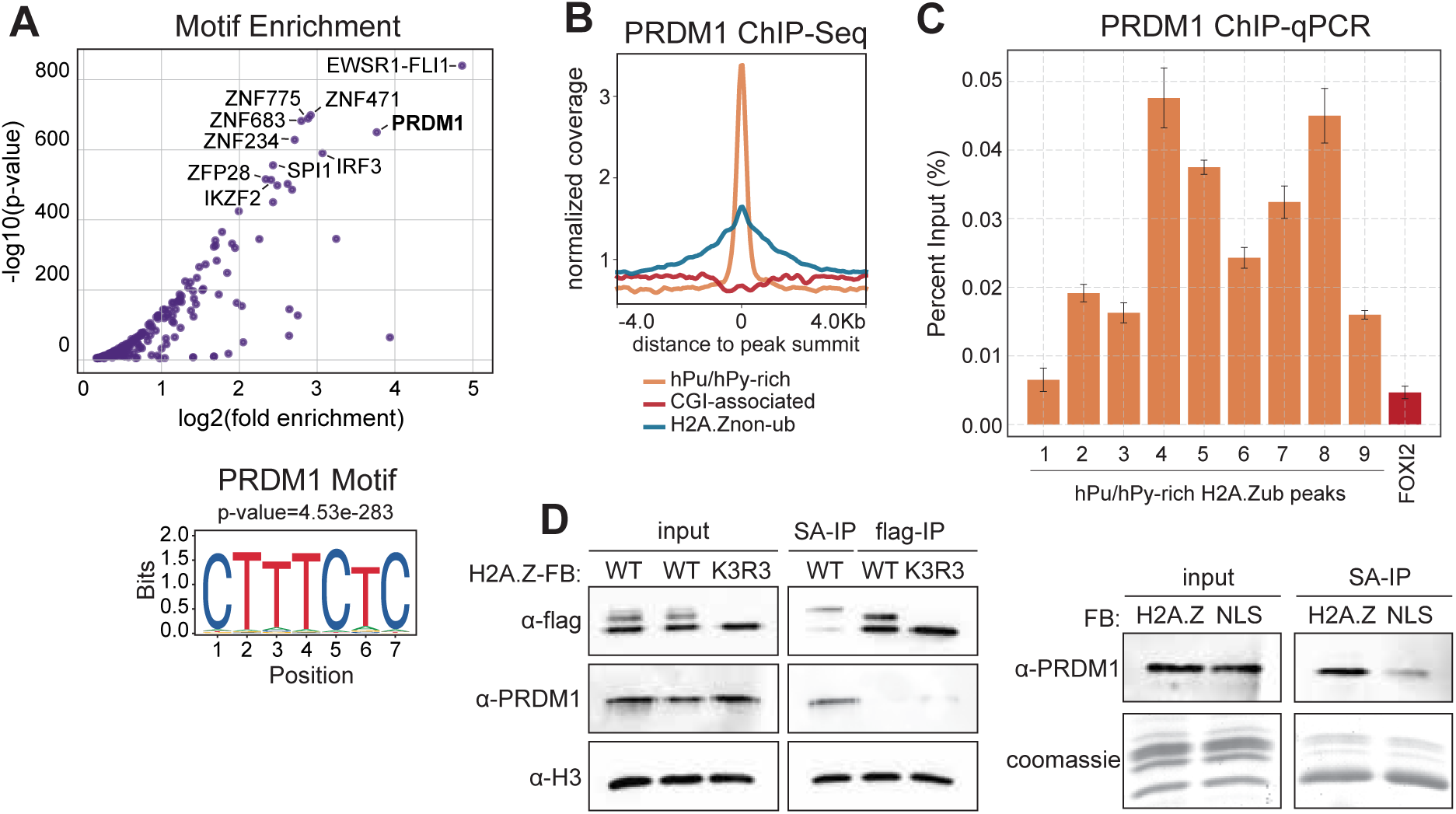
PRDM1 and H2AZub interaction at homopurine-homopyrimidine (hPu/hPy) tracts. **A.** Left: motif enrichment analysis of hPu/hPy-rich H2A.Zub peaks performed using SEA (MEME Suite). Total H2A.Z peaks were used as the input control. Right: position weight matrix of enriched PRDM1 motif. **B.** Density profile for PRDM1 ChIP-Seq signal at hPu/hPy-rich or CGI-associated H2A.Zub peaks and H2A.Znon-ub peaks. **C.** PRDM1 ChIP-Seq validation at hPu/hPy-rich H2A.Zub peaks from Fig. 6C. H2A.Zub peak with no hPu/hPy enrichment (FOXI2) serves as negative control. **D.** Left: PRDM1 co-immunoprecipitation in streptavidin (SA) and Flag-IP were examined by Western blot. Right: PRDM1 co-immunoprecipitation in H2A.Z versus NLS-FB SA-IP.

To explore a potential biochemical interaction between PRDM1 and H2A.Zub, we isolated H2A.Zub using our Ubiquitylation-dependent Auto-biotinylation (UAB) system and tested for PRDM1 co-immunoprecipitation (Fig. 7D). Due to low endogenous protein abundance, PRDM1 detection by Western blot was only possible following ectopic expression. Nonetheless, our results show that PRDM1 co-immunoprecipitated preferentially with H2A.Zub (SA-IP) compared to total H2A.Z (WT Flag-IP) or H2A.Znon-ub (K3R3 Flag-IP). Furthermore, PRDM1 co-immunoprecipitation was significantly higher in streptavidin pulldown from H2A.Z-FB + Avi-ub-expressing cells compared to NLS-FB + Avi-ub-expressing cells, indicating that enhanced PRDM1 recovery was attributable to H2A.Zub rather than non-specific binding to streptavidin beads or to biotinylated endogenous histones. Overall, these results provide evidence for co-localization of H2A.Zub and PRDM1 at hPu/hPy tracts and for a preferential biochemical interaction between the two.

## DISCUSSION

In this study, we devised a novel technique designed to selectively purify ubiquitylated H2A.Z (H2A.Zub) nucleosomes from mammalian cells for both biochemical and genomic analyses. To our knowledge, this is the first use of the BirA-Avi-tag biotinylation system to target and biotinylate a post-translational modification on a modified protein. The auto-biotinylation of Avi-ubiquitylated H2A.Z-FB is predicted based on the close proximity of the BirA and the Avi-tag once the Avi-ubiquitin is added to the C-terminus of H2A.Z-FB. One caveat of this system is that the H2A.Z-BirA fusion could theoretically reach and biotinylate other proximal proteins that were also Avi-ubiquitylated. To that end, in our nucleosome IP-Western blots (Fig. 1C), we did detect some biotinylation of a ∼25 kD protein that we believe to be the Avi-ubiquitylated endogenous H2A.Z. Nevertheless, our studies also showed the predominant biotinylated molecule in cell lysates was the Avi-ubiquitylated H2A.Z-FB, and the results from additional analyses matched well with the hypothesized functions of H2A.Zub (5,22,24). With the success of this proof of principle study, it is possible that a similar system can be adapted to purify and study other ubiquitylated proteins as well as other polypeptide post-translational modifications such as sumoylation.

Using our H2A.Z-UAB system, we provided direct evidence for the co-existence of multiple repressive histone modification hallmarks on H2A.Zub nucleosomes including high levels of H3K27me3, and low levels of H3K4me2/3 and H3K27ac. Conversely, non-ubiquitylated H2A.Z (H2A.Znon-ub) nucleosomes were enriched for active histone marks. These findings support the broader hypothesis that combinatorial histone post-translational modifications (PTMs) act in concert to mediate downstream regulatory effects (50). Furthermore, we report a ubiquitylation-dependent association of H2A.Z with DNA-methylated CpG islands and with unique homopurine-homopyrimidine sequences. Together, these discoveries add new insights into the PTM-dependent functional diversification of H2A.Z in transcriptional regulation, and the broad association of H2A.Zub with multiple types of silenced genomic domains.

Our ChIP-seq analyses provide the first genome-wide map of H2A.Zub enrichment and distribution within the human genome (Fig. 3). Since only a fraction of total H2A.Z in 293T cells is monoubiquitylated, we initially hypothesized that this subpopulation would be clustered at a subset of repressed H2A.Z-associated genes. However, we instead found that ubiquitylated and non-ubiquitylated H2A.Z are extensively intermingled throughout the genome. Furthermore, we observed comparable H2A.Zub levels at the TSSs of transcriptionally active and inactive genes, whereas total H2A.Z was significantly more abundant at the TSSs of active genes (Fig. 3A). This suggests that, while H2A.Zub levels remain relatively constant across gene classes, the ratio of non-ubiquitylated to ubiquitylated H2A.Z is higher at active promoters. This enrichment of non-ubiquitylated H2A.Z at active genes is consistent with its well-established association with transcriptional activity in human cells (4,17,51,52). Furthermore, the similar H2A.Z levels at the TSSs of low- and high-expressing genes are consistent with the model that H2A.Z functions to poise genes for activation, rather than directly influencing transcriptional output (Fig. 3A) (4,5,53). Altogether, our ChIP-seq data suggest that transcriptional activity does not correlate with the exclusive presence of either non-ubiquitylated or ubiquitylated H2A.Z but is more likely determined by the relative ratio of these two forms at promoters. In support of this idea, differentially enriched H2A.Zub and H2A.Znon-ub ChIP-seq peaks revealed clear correlations: promoter enrichment of H2A.Zub was associated with gene-silencing, while enrichment of non-ubiquitylated H2A.Z correlated with gene activation (Fig. 4B-D). More interestingly, we found that H2A.Zub peaks were not only hypermethylated for H3K27me3 but also hypomethylated for H3K4me3, whereas the opposite trend is observed at H2A.Znon-ub peaks (Fig. 4B). These trends exactly mirror the patterns of histone modification identified from our Western blot analyses of purified H2A.Zub nucleosomes (Fig. 4A). Therefore, our study provides corroborating biochemical and genomics evidence indicating reciprocal correlations between H2A.Z ubiquitylation and H3K4-/H3K27-methylation levels.

Previous ChIP-seq analyses of H2A.Z localization in mouse and human ES cells have identified a strong correlation/co-enrichment of H2A.Z and H3K4me3, especially at promoters and enhancers (4,5,53). Moreover, knockdown of H2A.Z or MLL4 in mouse ES cells has been reported to respectively cause reduced H3K4me3 or H2A.Z levels at some enhancers, suggesting that deposition of these epigenetic marks is functionally linked in some circumstances (4). In contrast, our studies reveal that there is also a striking anti-correlation between H2A.Zub and H3K4me3 (Fig. 4A-B), thus raising the possibility that ubiquitylation of H2A.Z antagonizes the connection between H2A.Z and H3K4 methylation or promotes H3K4me3 de-methylation.

H2A.Z and DNA methylation are recognized as opposing players in chromatin regulation; their genomic distributions show minimal overlap and the depletion of one leads to increased levels of the other (40,41,54,55). Our findings reveal that this antagonism applies only to non-ubiquitylated H2A.Z, whereas H2A.Zub displays a pronounced association with DNA methylation (Fig. 5A, F). Intriguingly, a significant portion of highly methylated H2A.Zub peaks correspond to CpG-rich islands (CGIs; Fig. 5D), and even more remarkably, nearly all H2A.Zub-enriched CGIs are methylated. In mammals, most CGIs remain unmethylated, with only a subset undergoing methylation in somatic cells (48). DNA hypermethylation at CGIs is a characteristic feature of human neoplasia, with certain cancers being classified as exhibiting a CGI methylator phenotype (CIMP) (56). Of note, HEK293 cells display a CIMP phenotype, with a higher number of methylated CGIs (28). When ranking CGIs based on their DNA methylation levels in HEK293T cells, we again see a significant portion of CGIs exhibiting high DNA methylation levels, consistent with the CIMP classification of this cell line (Fig. 5E). Importantly, we observe a pronounced enrichment of SA-/Flag-IP (H2A.Zub) signal at methylated CGIs and depletion at unmethylated sites, suggesting a robust association between H2A.Zub and DNA methylation (Fig. 5E). Indeed, there is a growing consensus regarding the pivotal role of ubiquitylated histones in both the establishment and maintenance of DNA methylation in human cells. For example, previous studies have highlighted the role of Polycomb-mediated H3K27me3 in marking CGIs for de novo methylation in cancer cells and during neuronal differentiation (57–59). More recently, it has been revealed that H2AK119ub, not H3K27me3, recruits DNMT3A to Polycomb-regulated CGIs, facilitating their methylation in both cancer cells and during normal development (42,44). H2AK119ub has also been implicated in recruitment of DNMT1 for maintenance of DNA methylation after replication (60). Given the structural similarities between H2AK119ub and H2A.Zub nucleosomes, it is tempting to speculate that H2A.Zub is also capable of recruiting DNA methyltransferases and may have a role in targeting DNMTs to selected CGIs for methylation.

Our current comprehension of the biological implications of homopurine-homopyrimidine (hPu/hPy) sequences remains very limited. Such sequences are found in mammalian genomes in much greater frequencies than predicted by chance and are often found near or upstream of transcriptional units (61,62). Our observed enrichment of H2A.Zub at these sequences is consistent with early studies that found correlation between hPu/hPy sequences and transcriptional repression (63). Whether or how these DNA sequences participate in transcriptional regulation is unclear. Some hPu/hPy sequences have mirror repeats that can form intramolecular triple DNA structures known as H-DNAs, and genome-wide evidence for their existence *in vivo* has only recently emerged and remains scarce (26,64). While the evolutionary benefits of these structures are unclear, it is evident that they can interfere with DNA-templated processes involving unwinding, such as replication and transcription (65). Moreover, H-DNA triplexes have been shown to promote homologous recombination and are notably concentrated at translocation hotspots (66). Consequently, H-DNA has been implicated in inducing point mutations and double-strand breaks (DSBs) (67). To date, the epigenetic features at hPu/hPy-tracts have not been thoroughly investigated. In this study, we demonstrate that these sequences exhibit an enrichment for ubiquitylated H2A.Z and H3K27me3 (Fig. 6). A notably high proportion of H2A.Zub peaks co-occupied by H3K27me3 overlap with hPu/hPy-tracts (Fig. 6B-D). Furthermore, enrichment for both H2A.Zub and H3K27me3 appears to be a characteristic of hPu/hPy-tracts genome-wide (Fig. 6E). Finally, we identify PRDM1, a repressive transcription factor implicated in the recruitment of Polycomb repressive complexes to chromatin, as being enriched at hPu/hPy-rich H2A.Zub peaks and biochemically associated with H2A.Zub-containing nucleosomes (Fig. 7). Together, these results support a model in which H2A.Zub enrichment at hPu/hPy-tracts contributes to the establishment of a chromatin environment that is less permissive to transcription, potentially mitigating DNA damage at these regions.

## ACKNOWLEDGEMENTS

We thank Keyur Advaryu and Kashif Aziz Khan for scientific discussions and general support, and Emanuel Rosonina for critical reading of the manuscript.

## AUTHOR CONTRIBUTIONS

Bakhtiyar Taghizada: Conceptualization, Data curation, Formal analysis, Investigation, Methodology, Software, Validation, Visualization, Writing—original draft, Writing—review & editing. Marlee K. Ng: Conceptualization, Data curation, Investigation, Methodology, Validation, Visualization, Writing—original draft. Ulrich Braunschweig: Formal analysis, Software, Visualization. Benjamin J. Blencowe: Project administration, Resources, Supervision. Peter Cheung: Conceptualization, Funding acquisition, Investigation, Project Administration, Supervision, Writing—original draft, Writing—review & editing.

## SUPPLEMENTARY DATA

Supplementary Data are available at NAR online.

## DATA AVAILABILITY

Raw and processed data generated in this study have been deposited in the Gene Expression Omnibus (GEO) under accession number GSE322838.

